# The Impact of Gag Non-Cleavage Site Mutations on HIV-1 Viral Fitness from Integrative Modelling and Simulations

**DOI:** 10.1101/2020.07.05.188326

**Authors:** Firdaus Samsudin, Samuel Ken-En Gan, Peter J. Bond

## Abstract

The high mutation rate in retroviruses is one of the leading causes of drug resistance. In human immunodeficiency virus type-1 (HIV-1), synergistic mutations in its protease and the protease substrate – the Group-specific antigen (Gag) polyprotein – work together to confer drug resistance against protease inhibitors and compensate the mutations affecting viral fitness. Some Gag mutations can restore Gag-protease binding, yet most Gag-protease correlated mutations occur outside of the Gag cleavage site. To investigate the molecular basis for this, we now report multiscale modelling approaches to investigate various sequentially cleaved Gag products in the context of clinically relevant mutations that occur outside of the cleavage sites, including simulations of the largest Gag proteolytic product in its viral membrane-bound state. We found that some mutations, such as G123E and H219Q, involve direct interaction with cleavage site residues to influence their local environment, while certain mutations in the matrix domain lead to the enrichment of lipids important for Gag targeting and assembly. Collectively, our results reveal why non-cleavage site mutations have far-reaching implications outside of Gag proteolysis, with important consequences for drugging Gag maturation intermediates and tackling protease inhibitor resistance.

## Introduction

RNA viruses are among the most adaptable pathogens threatening human health. With their high mutation rates^1^, they are orders of magnitude more adaptable than most DNA-based organisms^2^, resulting in their ability not only to escape the immune system and become drug resistant, but jump across host species boundaries causing emerging infectious diseases that could turn into global pandemics^3-6^. HIV-1 represents one of the most extensively characterized RNA viruses, with mathematical modelling showing that many clinically established drug resistance mutations could occur within the first replication cycle^7^, and every single possible point mutation could occur thousands of times within a single day, leading to rapid drug resistance^8^, hence leading to the need to treat HIV patients with a cocktail of multiple antiviral drugs with different mechanisms^9^. Understanding the role of these mutations is thus an imperative aspect in novel antiviral drug development.

Most clinically approved HIV treatments target various key enzymes that are critical to the life cycle of the virus, one of which is the HIV-1 protease. HIV-1 protease cleaves the structural polyproteins Gag and Gag-Pol into their mature components to create the infectious virion. To inhibit this, protease inhibitors (PIs) bind competitively to the viral protease to prevent Gag substrate proteolysis and virion maturation. PIs form a vital second line component of the antiretroviral therapy (ART) in the management of adults with HIV. As failure of first line regimens develop, a potent second line drug is crucial.^10^ This is often associated with multi nucleoside reverse transcriptase inhibitors (NRTIs) resistance. Ritonavir-boosted PIs represents the only available second-line treatment option for the majority of HIV patients worldwide; however, emerging mutations in HIV-1 protease often render PIs ineffective, with some mutations even leading to cross-drug resistance^11-13^. This is further aggravated by synergistic mutations on the Gag polyprotein itself^14-17^, which is the rationale for elucidating the drug resistance encoded in Gag. Gag mutations around the protease cleavage site have been shown to restore Gag-protease binding by introducing new chemical interactions and inducing subtle conformational changes that compensate for the loss of affinity due to the mutations on HIV-1 protease^18^. Apart from mutations around the protease cleavage sites, Gag also harbours mutations away from these regions, many of which have been shown to directly contribute towards PI resistance^19-21^. To date, the role of these non-cleavage site mutations is largely unknown, with some analysis pointing to allosteric communication^22^. A study using paramagnetic NMR spectroscopy showed that Gag can form transient complexes with protease and the binding interface involves non-cleavage site regions that are prone to mutation causing drug resistance^23^. In this study, we aim to use structural modelling and simulations to decipher how Gag non-cleavage site mutations contribute towards the overall fitness of the HIV-1 virus.

The HIV-1 Gag polyprotein is a 500 amino acid precursor protein containing the matrix (MA), capsid (CA), nucleocapsid (NC) and P6 domains, as well as two spacer peptides, SP1 and SP2. During the late phase of the HIV-1 replication cycle, Gag is sequentially cleaved by the protease enzyme into these domains (Figure S1), which subsequently form an infectious mature virion^24^. Each domain plays a specific role during maturation; for example, the MA domain drives full-length Gag to assemble at the plasma membrane of the host cell, while the CA and NC domains encapsulate the viral RNA genome. The CA domain also interacts with CypA, which is a host cell protein that is incorporated into the virion and is essential for capsid uncoating^25,26^. Due to its functional significance in the HIV-1 life cycle, Gag is an attractive target for therapeutic agents and to date, several drugs that target the CA domain have been identified^27^. Nevertheless, a Gag inhibitor is yet to be clinically approved. For example, Bevirimat is a drug candidate that stabilises the immature Gag lattice by preventing proteolysis between the CA and SP1 domains^28^. A clinical trial, however, showed a reduced response amongst certain patients due to the highly polymorphic nature of Gag across different circulating HIV-1 subtypes and various resistant mutations^29,30^, which highlights the importance of understanding the role of Gag non-cleavage site mutations.

Over the past few years, integrative modelling and molecular dynamics (MD) simulations, in tandem with advances in structural biology, have provided valuable insights into the molecular mechanism underlying viral function and dynamics^31^. In the study of HIV-1, this ranges from multiscale simulations of the entire viral capsid shell and its assembly pathways^32-37^, to simulations of individual Gag proteins with host cell components, such as the CA domain with CypA^38^, kinesin-1 adaptor protein FEZ1^39^ and inositol hexakisphosphate (IP6)^40^, as well as the MA domain with a model plasma membrane^41^. A previous structural model of full-length monomeric HIV-1 Gag revealed allosteric communications between non-cleavage site mutations and the first Gag cleavage site^22^, providing a glimpse into how residues far away from the protease cleavage site could affect proteolysis. However, this model did not take into account *in vivo* Gag oligomerisation, which is instrumental during virus particle maturation. Crystal structures show that the MA domain exists as a trimer^42^, whilst the CA domain forms a hexamer^43^. An electron microscopy (EM) study of the MA protein in a PIP2 containing membrane showed that under higher order conditions, it organizes into hexamers-of-trimers^44^.

Structural models of sequentially cleaved Gag are imperative for understanding the conformational changes involved upon proteolysis, which may improve our overall knowledge of Gag mutations and potential drugging of these intermediates. Using available structures of HIV-1 Gag domains, we built an integrative model of the complete oligomeric Gag polyprotein cleavage product (Figure 1) bound to a viral membrane model (Figure 2). Based on coarse grained (CG) MD simulations using the Martini forcefield of the wild-type (WT) and mutant Gag variants, supported by careful calibration against atomic-resolution sampling, we found that non-cleavage site mutations can interact with cleavage site residues and potentially alter their local environment, while mutations on the MA domain confer stronger binding to the plasma membrane. Subsequent all-atom MD simulations of the CA domain revealed how mutations in this region modulate CypA interaction as well as stabilise oligomer formation. Overall, our study uncovers how these distant mutations can affect various processes during HIV-1 viral maturation and contribute towards its overall fitness.

**Figure 1:**
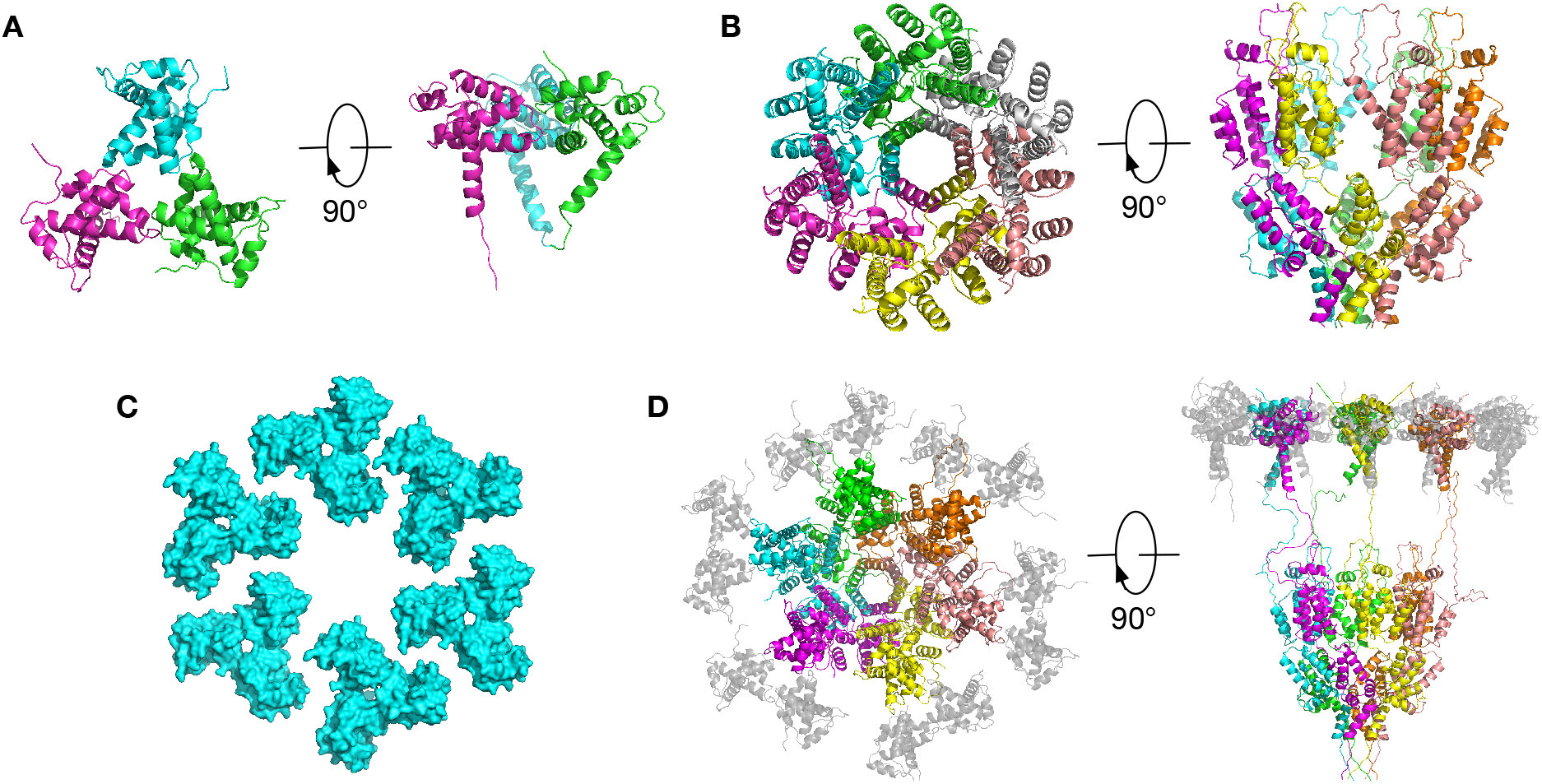
Modelling HIV-1 Gag MA-CA-SP1. (A) The crystal structure of the MA trimer (PDB: 1HIW). (B) The crystal structure of the immature CA hexamer (PDB: 5L93). (C) The hexamer of trimers model for the MA domains as supported by EM. (D) A model of the MA-CA-SP1 hexamer. The central subunits of the MA trimers are connected to the CA domains and coloured as in (B), whereas the peripheral MA subunits are coloured grey.

## Results

### Membrane dynamics of the HIV-1 Gag MA-CA-SP1 polyprotein

The Gag polyprotein is sequentially cleaved by HIV-1 protease during viral maturation (Figure S1). It is unknown which state of viral maturation is affected by non-cleavage site mutations. As such, we aim to investigate the effects of these mutations on both immature and mature states of Gag, where structures are available. The first cleavage occurs between the SP1 and NC domains to produce the MA-CA-SP1 polyprotein, which represents the largest Gag cleavage product. A hexameric model of the HIV-1 Gag MA-CA-SP1 polyprotein was built using the crystal structures of the MA trimer^42^ and CA hexamer^43^ (see Methods section for further details) (Figure 1). The crystal structure of the immature CA hexamer was aligned with six copies of MA trimers (hexamers-of-trimers) following the arrangement observed in membrane bilayers^44^. A solution NMR structure of an N-terminal fragment of immature HIV-1 Gag that includes MA and the C-terminal portion of CA shows that the linker between MA and CA domains is inherently flexible and adopts multiple conformation^45^. This is supported by another NMR study of an HIV-1 Gag fragment consisting of the MA, CA, SP1 and NC domains that showed individual domains orient in a manner semi-independent of each other, with adjoining linkers that are intrinsically disordered^46^. As such, we constructed a flexible intervening linker between the MA and CA domains *ab initio*. Gatanaga et al. discovered PI-resistant Gag variants with non-cleavage site mutations mapped to the MA-CA-SP1 complex (Table S1, Figure S1)^20^. These include seven unique mutations, with five located in the MA domain and two in the CA domain. We generated models for three variants (specifically variant numbers 3, 6, and 7, as outlined in Table S1) that cover all of these seven unique mutations (Figure 2A). These models were built by substituting the wild-type residues with the corresponding mutant version, assuming that these point mutations do not have any large structural impact upon Gag. To understand the conformational dynamics of the polyprotein and the potential roles of these non-cleavage site mutations, we performed four independent 15 μs CG MD simulations of the MA-CA-SP1 model bound to a realistic HIV-1 membrane model (Simulations 1-4 in Table S2). The membrane model contained phosphatidylcholine (PC), phosphatidyl ethanolamine (PE) and phosphatidylserine (PS) lipids, as well as PIP2, sphingomyelin and cholesterol, as determined by a previous lipidomics study^47^ (see Methods section for further details), while the MA domain N-terminus was myristoylated (Figure 2B).

As previously predicted^41^, the myristate group on the N-terminal glycine residue of MA domain effectively anchored the protein to the membrane. Within the first few nanoseconds, the linker connecting the MA and CA domains contracted, effectively pulling the CA-SP1 domain towards the membrane (Figure 2B). To verify the MA-CA linker contraction observed during CG sampling, we also performed four independent 500 ns atomistic simulations of the MA-CA-SP1 monomer with positional restraints applied to the backbone atoms of lipid binding residues to mimic membrane association (Simulation 5 in Table S2). Additionally, we conducted CG simulations of the same monomeric system (Simulation 6 in Table S2). Both of these additional sets of simulations showed a contraction of the MA-CA linker, resulting in a similar distribution of distances between the MA and CA domains compared to the CG hexamer simulations (Figure S2). While the atomistic simulations initially sampled a wider spread of distances, after around 300 ns this converged to 4-6 nm, reproducing the CG distributions. This validates our CG hexamer model and shows that the linker region does not maintain an extended conformation, but rather contracts to allow direct contact between the MA and CA domains.

Curvature has been shown to be important in modulating the interface of the CA lattice^43^,^48^. Structural flexibility of the CA N-terminal and C-terminal domains give rise to a curved CA-CA interface, and oligomerisation of this curved interface at the plasma membrane of infected cells results in membrane bending and subsequent budding of the virus particles. The structure of the CA hexamer used in our modelling was taken from the intrinsically curved CA lattice within immature virus-like particles^43^. However, the MA domains were arranged to be flat to match the flat model membrane at the beginning of the simulations (Figure 1D). Interestingly, over the course of the simulations, as the MA-CA linker contracted allowing direct contacts between these two domains, the MA domain spontaneously adapted to the curved CA hexamer (Figure 2B and S3). Consequently, the curved MA domain induced some degree of curvature in the membrane. This shows how a curved CA-CA interface could affect surrounding Gag domains and the membrane, thereby affecting viral release.

**Figure 2:**
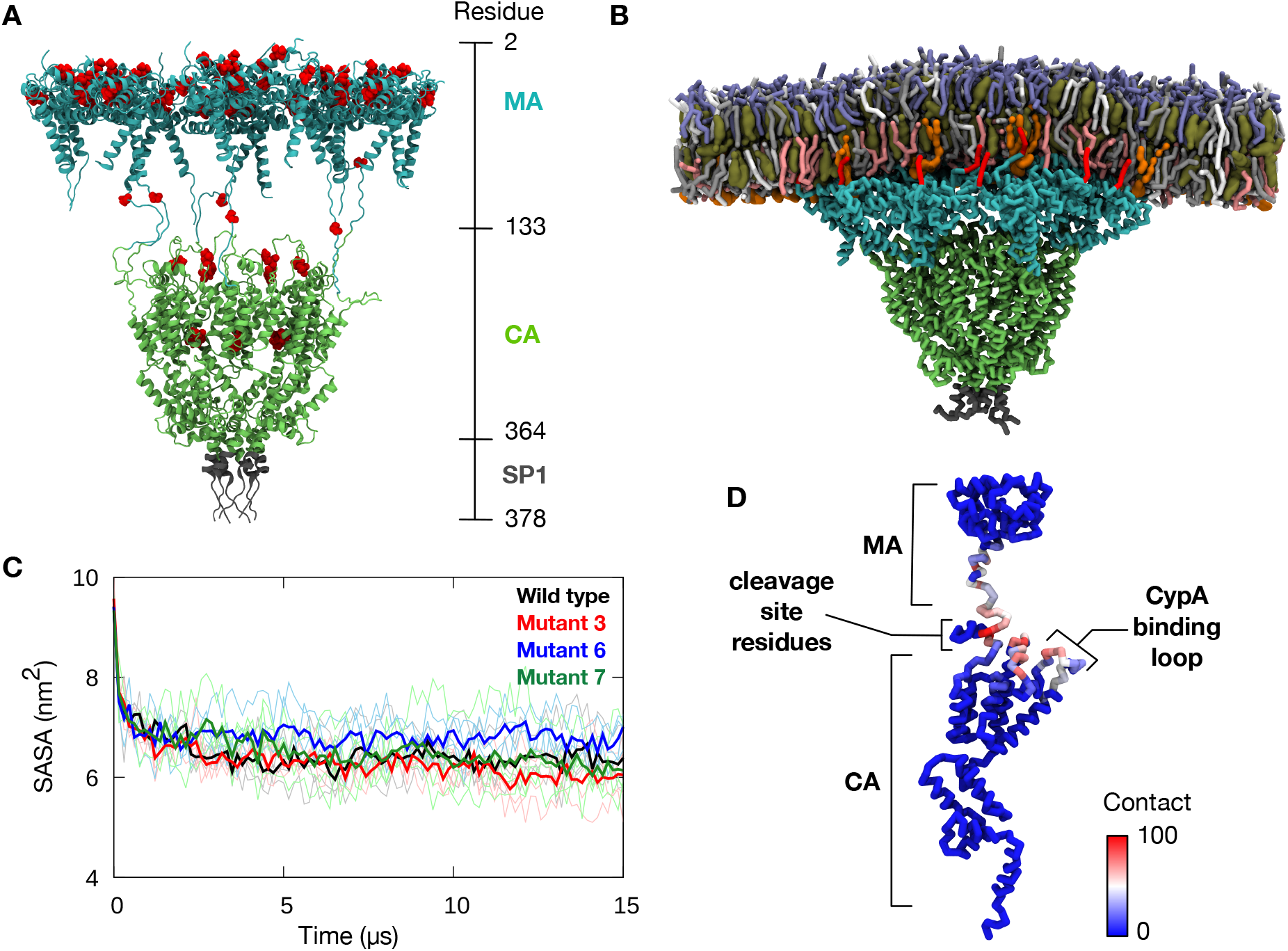
HIV-1 Gag MA-CA-SP1 model. (A) An atomic model of MA-CA-SP1 hexamer built using the crystal structures of MA trimer^42^ and immature CA hexamer^43^, shown in cartoon representation. The MA domain (cyan), CA domain (green), and SP1 domain (dark grey) are shown in different colours. The positions of seven unique non-cleavage site mutations^20^ are shown in red. Residue numbering for each domain and their approximate positions are shown on the right. (B) The final snapshot from one of the 15 μs CG simulations of a membrane-bound WT MA-CA-SP1 hexamer showing contraction of the linker connecting MA and CA domains. Protein backbone and lipids are shown in licorice representation. N-terminal myristate (red), PC (white), PE (light grey), PS (pink), PIP2 (orange), sphingomyelin (purple) and cholesterol (dark green) are shown in different colours. (C) Solvent accessible surface area (SASA) of the MA-CA cleavage site region (residue 128-137) over the course of 15 μs CG simulations. Thin lines represent the six individual subunits and thick lines show the running average. Data were averaged over four independent simulations. (D) Average percentage of contact made by the cleavage site residues with the rest of the protein, mapped onto the backbone of a single subunit of MA-CA-SP1 from the WT simulations. Data were averaged over six subunits and four independent simulations. The cut-off distance used for contact analysis was 0.6 nm.

### Interaction between cleavage site residues and non-cleavage site mutations

The linker contraction described above resulted in a decrease in solvent accessibility of the cleavage site residues between the MA and CA domains from ~10 nm^2^ to ~6 nm^2^ (Figure 2C and S4). In the first round of proteolysis, Gag is cleaved at the SP1/NC cleavage site to produce MA-CA-SP1 and NC-SP2-P6 intermediates (Figure S1). In the second round of proteolysis, P6 is cleaved off the latter at a rate measured in vitro approximately 9-fold slower than the initial cleavage, while MA is cleaved off the former at an estimated 14-fold slower rate than the first round of proteolysis^49^. Cleavage at the MA/CA site therefore happens around 1.6 times more slowly than at the SP2/P6 site. Our simulations suggest that this may be caused by the reduction in exposure of the MA/CA cleavage site, whereby at the end of the simulations, the solvent accessible surface area (SASA) of the cleavage site is also approximately 1.6 times smaller than at the beginning of the simulations, due to the contraction of the linker region between MA and CA domains. This suggests that the MA-CA linker dynamics could account for the difference in the rates observed in Gag cleavage, and that the SP2-P6 linker adopts a more extended and less buried conformation.

This linker contraction also led to the cleavage site residues interacting with neighbouring subdomains, predominantly with the C-terminal helix of the MA domain, N-terminal loop of the CA domain, and several inter-helical loops on the CA domain, including the CypA binding loop (Figure 2D). Two PI-resistant mutations were identified at the interaction sites – G123E and H219Q. To understand how these mutations may affect the cleavage site residues, contact analysis with a cut-off distance of 0.6 nm was performed between the cleavage site residues and these two specific positions in the WT and mutant proteins, particularly focusing on potential inter- and intra-subunit interactions. Due to their close proximity, we found that the residue at position 123 made contact primarily with the N-terminal portion of the cleavage site from the same Gag subunit (Figure 3A and 3B). Both WT (glycine) and mutant (glutamate) residues showed a similar percentage of contact over the course of the simulations. “Back-mapping” of CG simulation snapshots to transform them into atomic resolution, performed using a flexible geometric approach^50^ (details in Methods section), suggested that a glutamate residue at this position may interact with polar residues on the cleavage site, such as Q130 and Y132 (Figure 3C). We performed three independent 200 ns atomistic simulations of a single MA-CA-SP1 subunit to refine the back-mapped structure, and found that the glutamate residue indeed interacted primarily with Y132 (Figure S5). Additionally, the residue also contacted residues N131 and Q130. In contrast, a glycine residue at this position interacted primarily with V128 and S129. Given the change in the overall size and charge of the residue in the WT and mutants (from small and neutral to large and acidic), the G123E mutation alters the accessibility and electrostatic properties in the vicinity of the cleavage site and would therefore be expected to directly interfere with proteolysis.

While the interactions of G123E with the cleavage site residues occur within the same subunit of the MA-CA-SP1 polyprotein, inter-subunit interactions were observed for H219, whereby this residue contacted the cleavage site of an adjacent subunit (Figure 3D). The frequency of such interactions, however, was notably lower than that displayed by G123E, and interestingly, the mutation to glutamine resulted in a reduced contact with the cleavage site (Figure 3E). Back-mapping to atomistic representation suggested potential for interactions with polar residues such as N131 and N137 (Figure 3F). Atomistic simulations of two adjacent MA-CA-SP1 subunits using the back-mapped structure showed that histidine at this position made intermittent contacts with I134, V135, Q136 and N137, whilst glutamine interacted primarily with Y132 (Figure S5). Given that the WT histidine may potentially become protonated, it may affect local electrostatic surface properties of the Gag cleavage site to modulate the catalytic efficiency of the protease enzyme.

Overall, our data suggest that even though these mutations occur far from the protease binding site, they can physically interact with the cleavage site residues, either within the same Gag subunit or with a neighbouring subunit, as the linker region between MA and CA domain contracts following the first proteolytic cleavage.

**Figure 3:**
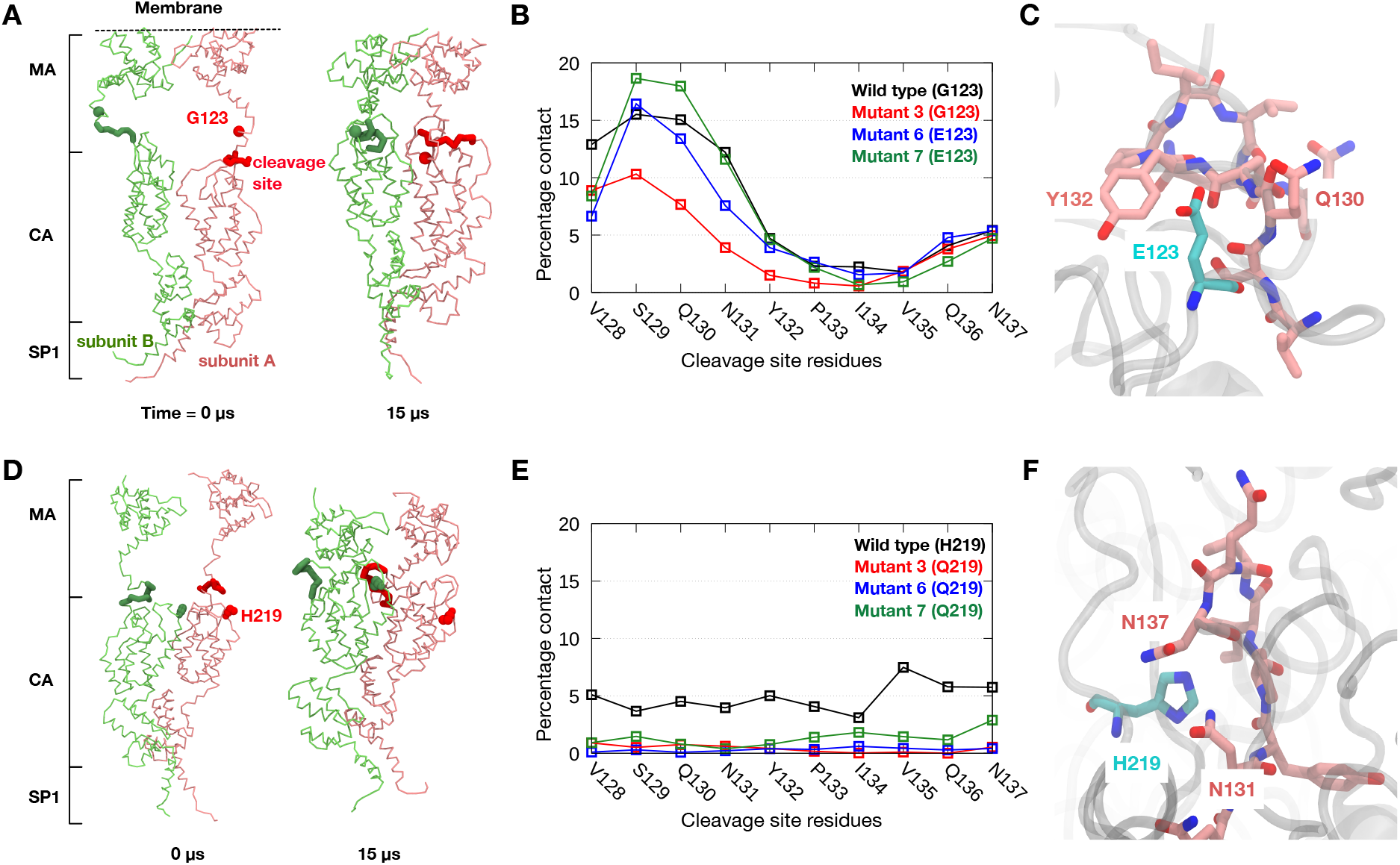
Interactions between non-cleavage site mutations and cleavage site residues. (A) Snapshot at the beginning and end of the CG simulations of WT MA-CA-SP1 showing the interaction of residue G123 (sphere representation) and cleavage site residues (thick licorice representation). Two subunits are shown in red and green. (B) Percentage of contacts across the entire CG simulation sampling made between G123/E123 with each of the residues of the MA-CA cleavage site. This is averaged over the six subunits of MA-CA-SP1 and the four independent trajectories. The cut-off distance used for contact analysis was 0.6 nm. (C) Atomistic model, derived from the final snapshot of one of the CG simulations, highlighting E123 in mutant Gag (cyan) and nearby cleavage site residues (pink). (D) Snapshot at the beginning and end of the CG simulations showing the interaction between H219 from one subunit to cleavage site residues on an adjacent subunit. (E) Contact analysis similar to (B) for H219/Q219. (F) Atomistic model derived from the CG simulations of WT Gag highlighting H219 in WT Gag (cyan) and surrounding cleavage site residues (pink).

### Mutations in the MA domain modulate membrane binding

Four of the mutations in the MA domain map onto the plasma membrane associating surface of our model, and three mutations – E12K, E40K and L75R – involve a switch to basic amino acids. We hypothesized that these mutations may therefore alter key electrostatic interactions with lipids. Indeed, in the three mutant variants, there is a notable increase in positive electrostatic charge on the membrane binding region compared to WT (Figure S6). In the WT Gag MA domain, the positive charge is concentrated on the peripheral region of the membrane binding surface of the MA trimer. In the mutant variants, in particular mutants 6 and 7, the positive charge covers almost the entire membrane binding surface including the central region at the interface of the MA trimer subunits.

Previous biochemical and structural studies showed that PIP2 is instrumental in targeting Gag to the plasma membrane of the host cell, and that it binds directly to the MA domain^51^,^52^. To understand how the changes in electrostatic surface properties caused by the PI-resistant mutations affect membrane binding, we determined the percentage of PIP2 interacting with MA throughout our CG simulations. This is based on the number of PIP2 molecules found within 0.6 nm of MA normalized by the total number of PIP2 molecules within the membrane. Over the course of the 15 μs simulations, there was a significant enrichment of PIP2 lipids around the MA domain, consistently observed across all trajectories, from around 30% at the beginning of the simulations to 60-80% by the end (Figure 4A). A number-density map averaged over the last 5 μs of the simulations showed a significantly higher density of PIP2 around the edges of MA hexamer-of-trimers as well as at the interface between MA trimers compared to areas further away from the protein (Figure 4B and S7A). This was due to electrostatic attraction between the anionic PIP2 headgroups and the positively charged membrane-peripheral surface of the MA domain. Interestingly, we did not observe any increment in the percentage of anionic PS lipids despite the higher concentration of PS compared to PIP2 in the membrane (Figure S7B and S7C). This may be due to the higher negative charge on the headgroups of PIP2 lipids. PIP2 enrichment around the MA domain is in agreement with previous experimental studies showing that PIP2 is responsible for anchoring HIV-1 Gag to the plasma membrane^53^. While both WT and mutant variants displayed PIP2 enrichment, all three mutants showed a larger degree of accumulation of PIP2 around MA. More importantly, the E40K and L75R mutations, which map around the interface of the MA trimer subunits, formed novel binding sites for PIP2 lipids (Figure 4C, 4D). Our simulations therefore suggest that these non-cleavage site mutations enhance interactions between the MA domain and PIP2 lipids, and may consequently improve membrane targeting of HIV-1 Gag during viral assembly.

**Figure 4:**
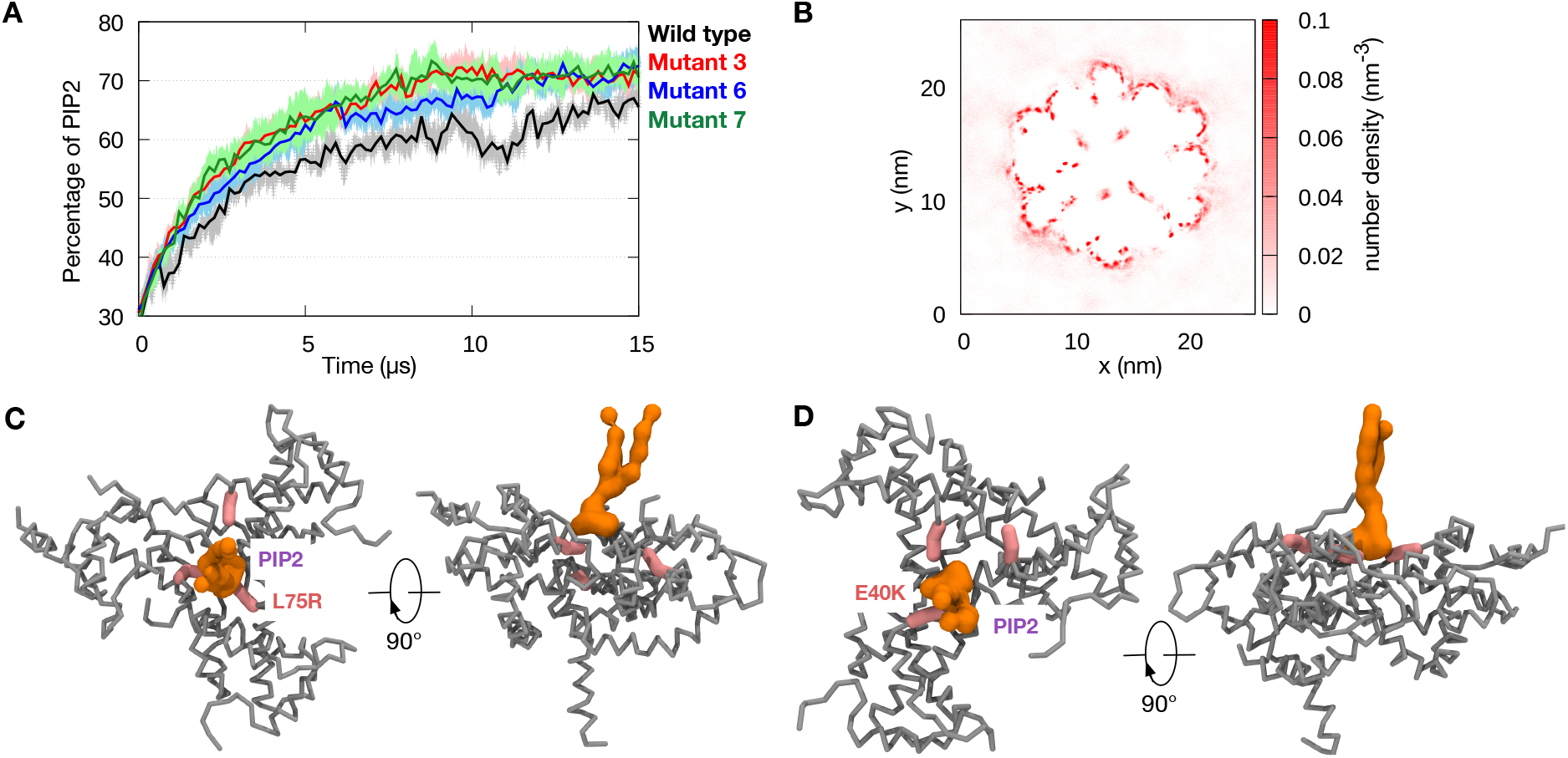
PIP2 is enriched around MA domain. (A) The percentage of PIP2 found within 0.6 nm of the MA domain during CG simulations of WT and mutant MA-CA-SP1 models. Error bars indicate standard deviation from four independent simulations. (B) Two-dimensional number density map of PIP2 for CG simulations of mutant 3 (Simulation 2 in Table S2) averaged over the last 5 μs. (C) Snapshots from one of the simulations of mutant 3 showing PIP2 binding at the interface of the MA trimer. The position of mutant residue, L75R, is shown in pink stick representation, whereas PIP2 is shown in orange surface representation. The rest of the MA domain is shown in grey. (D) Snapshot from simulations of mutant 7 (Simulation 4 in Table S2) showing PIP2 binding facilitated by E40K mutation, displayed in similar colours and representation as in (C).

### Mutations in CypA-binding loop optimize CypA interaction

During viral maturation, CA molecules undergo a large conformational change to form a conical capsid encapsulating the viral genome. We next sought to elucidate the impact of non-cleavage site mutations on the mature CA using the crystal structure of the mature CA hexamer^40^. We first investigated the interactions between mature CA and Cyclophilin A (CypA). CypA is a host protein that is recruited by HIV-1 Gag into mature virion particles. While the exact role of CypA in HIV-1 replication remains unclear, it is thought that the prolyl isomerase activity destabilises interactions between the subunits of the CA hexamer and therefore promotes viral core disassembly during the early stage of the replication cycle^26,54,55^. An optimal population of CypA in the infected target cell has been shown to be critical in modulating HIV-1 infectivity^56^. At a very low CypA concentration, the HIV-1 replication rate is severely reduced as the CA subunits are tightly bound to one another, which hinders virion uncoating. On the other hand, at a very high CypA concentration, interactions between CA subunits are greatly destabilised, resulting in an unstable virion core and thereby delayed virion maturation^57^. CypA binds to CA via an exposed proline-rich unstructured loop region in the N-terminal domain of the latter^55^. Interestingly, this loop houses H219Q, a mutation found in all seven variants from Gatanaga et al. (Table S1). To investigate the potential role of this mutation *vis-à-vis* CA-CypA interactions, we performed two independent 500 ns atomistic MD simulations of the WT and H219Q mutant of the mature CA hexamer either in its CypA-free (apo) or CypA-bound state (Simulations 7-12 in Table S2). The predicted pKa values of H219 in all six chains of the hexamer were determined using the PROPKA3 programme^58^ and were found to range between 6.3-6.5 (Table S3). As these pKa values are close to physiological pH, it is possible that H219 may exist in either protonated or unprotonated form. As such, the simulations of WT CA were performed with H219 in both protonation states. IP6 has been shown to be an important co-factor for HIV-1 capsid maturation and assembly^40,59-61^. In the immature Gag hexamer, IP6 interacts with K290 and K359, thus facilitating the formation of a six-helix bundle found between the CA and SP1 domains. Proteolytic cleavage between these two domains releases IP6 to subsequently bind to residue R18 in the mature CA, promoting assembly of mature CA hexamers and the lattice. As such, we included the IP6 molecule in all of our atomistic simulations of the mature CA hexamer (see Methods section for further details).

Our apo simulations revealed that the CypA binding loop was the most dynamic region of the entire CA molecules (Figure 5A). This is consistent with a recent structure of a CA tubular assembly determined by magic-angle spinning NMR by Manman et al. showing that the loop adopts at least four distinct conformations^62^. No noticeable difference was found between the flexibility of this loop in simulations with the neutral H219 *versus* the H219Q mutant (Figure 5B). Interestingly, when this histidine residue was protonated, we saw a slight increase in the root mean square fluctuation (RMSF) values of the loop, especially around G221-P222, which is the putative substrate for the CypA rotamase activity^63,64^. Cluster analysis performed on structures extracted from the simulations revealed that the CypA binding loop containing the protonated H219 was the most heterogenous with the largest distribution of root mean square deviation (RMSD) values, the highest number of distinct clusters, and the fewest structures grouped into the top three clusters (Figure S8). An increased association was observed between protonated H219 and E230, found on the opposite side of the loop (Figure 5C). Similarly, the central structure of the top cluster from protonated H219 simulations showed formation of a salt bridge between the protonated histidine and E230 (Figure 5D), which was absent from the top three clusters extracted from simulations with neutral H219 and H219Q mutant, as well as the ensemble of wild type CA structures solved by Manman et al.^62^ (Figure S8). However, this salt bridge was intermittent (Figure 5E), such that the CypA-binding loop may adopt alternative conformational states when H219 is protonated. This means that under low pH conditions, the transition between these two states increases the overall flexibility of the loop, which consequently may make it more difficult for CypA to bind. Mutation of H219 to glutamine would therefore abolish the ability of the loop to adopt these two conformational states, reducing the loop flexibility, and facilitate initial CypA binding. This would be advantageous inside the endosome which is associated with a lower local pH, to ensure sufficient CypA binding for core disassembly^65^.

**Figure 5:**
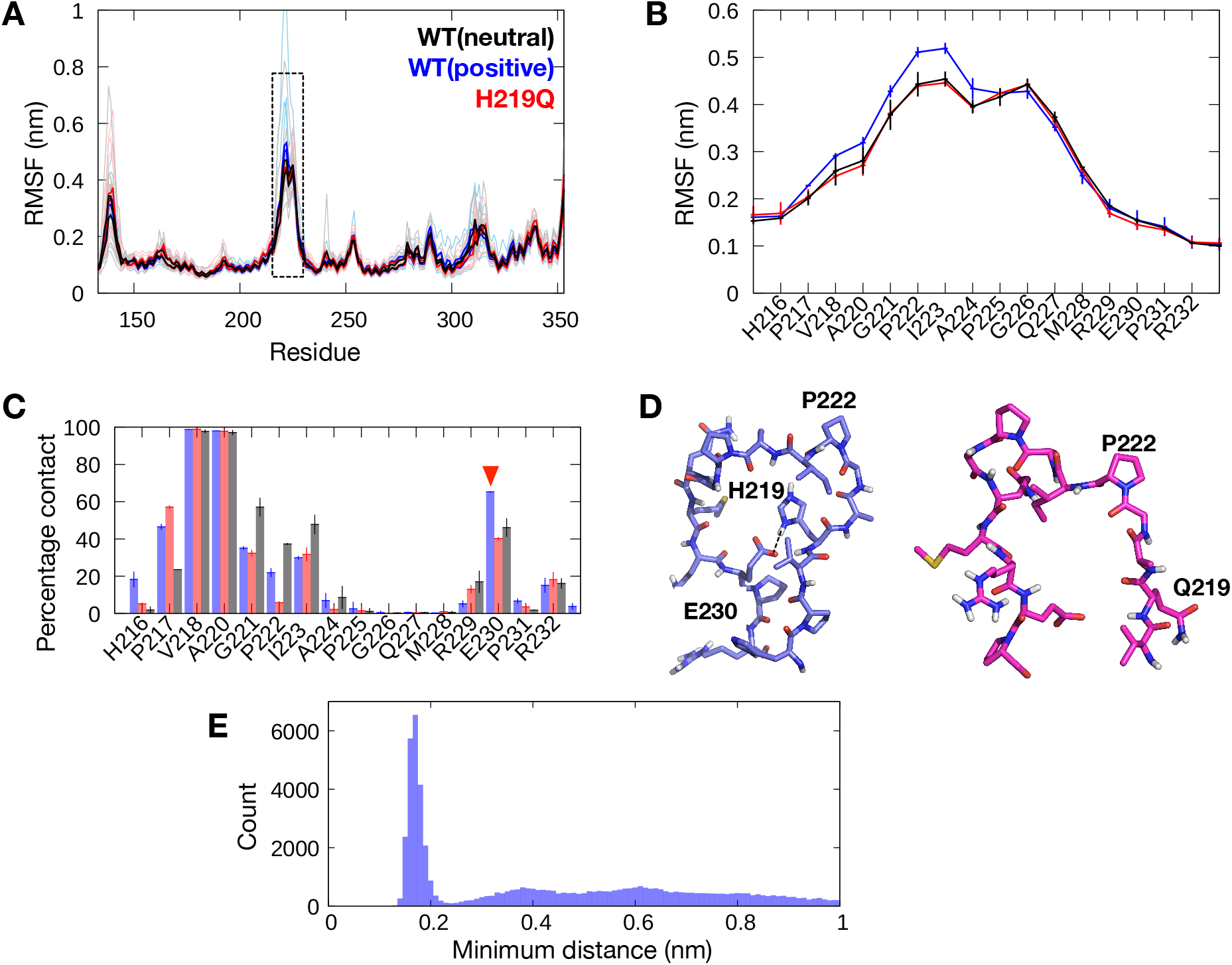
The dynamics of CypA-binding loop. (A) Per-residue root mean square fluctuations (RMSF) of the CA from two independent 500 ns apo simulations. Thin lines show the RMSF of each individual CA subunit, whilst the thick lines show the average from each simulation. Simulation with neutral H219 is shown in black, positively charged H219 in blue, and mutant H219Q in red. Dotted box indicates the loop region where CypA binds. (B) Enlarged RMSF plot for CypA-binding loop. Average from the two simulations is shown and error bars indicate standard deviations between repeat simulations. (C) Atomic contact analysis performed between residues at position 219 and the rest of the CypA-binding loop, including E230 (indicated by red arrowhead). Average values from two simulations are shown and error bars indicate standard deviations between repeat simulations. Cut-off distance for contact analysis is 0.4 nm. (D) Representative structures of the CypA-binding loop from the simulations with positively charged H219 (left) and mutant H219Q (right) calculated using cluster analysis. The central structures (the structure with the lowest RMSD from all other structures) of the top clusters are shown (more details in Figure S8). In the former, H219 can form a salt bridge interaction with residue E230 found on the opposite side of the loop, represented as dotted line. (E) The distribution of minimum distance between the hydrogen atoms bonded to the two nitrogen atoms on the side chain of the protonated H219 and the two oxygen atoms on the side chain of E230. Data taken from all six CA subunits and both repeat simulations.

Since the interactions between H219 and CypA form a part of the binding interface^55^, this raises the question of whether a mutation to glutamine would undermine the strength of binding. To investigate the potential effects of the H219Q mutation upon CypA binding affinity, we performed two independent 500 ns atomistic MD simulations of a mature CA hexamer with each of the subunits bound to a CypA molecule (Figure 6A). While the average buried surface area between CA and CypA in the WT and mutant CA systems were comparable throughout the simulations, we observed transient detachment of CypA molecules in two of the CA subunits from the H219Q variant (Figure 6B). Based on the crystal structure^55^, H219 forms hydrogen bonds with residues in the CypA active site, including N71. Our simulations revealed that this histidine residue also interacts with other polar residues such as T73, N102, and Q111 on CypA. In the H219Q mutant, the contacts made by the glutamine residue followed a largely similar pattern, although the frequency of interactions was noticeably lower compared to both neutral and protonated histidine (Figure 6C). Both of these analyses of CypA-bound CA simulations suggest that the H219Q mutation may potentially weaken CypA binding to the CA domain. This mutation is therefore likely beneficial in a CypA-rich environment whereby a high population of CypA in target cells may destabilise the viral core. Taken together, the H219Q mutation on the CypA-binding loop may play a crucial role in fine-tuning CypA interactions, especially in low endosomal pH and CypA-enriched infected target cells.

It is worth noting that cryo-EM studies of the large HIV-1 capsid lattice showed that a single CypA molecule could simultaneously bind two CA hexamers (Figure S9)^38,66^. To one hexamer, CypA binds to one CA subunit via canonical interactions with the CypA-binding loop discussed in this study. To the other neighbouring hexamer, CypA binds to two adjacent CA subunits via non-canonical interactions, facilitated by residues 254-257 as well as 227-229. We compared the RMSF values of these residues between our three sets of apo simulations involving neutral H219, protonated H219 and mutated H219Q variants (Simulations 7-9 in Table S2). There was no significant difference between the RMSF values for these residues, suggesting that the protonation and mutation of H219 would not affect the dynamics of residues involved in non-canonical binding of CypA. Interestingly, the cryo-EM structures show that these two non-canonical binding interfaces have a much smaller interaction area compared to the canonical interface, suggesting a weaker interaction. This is supported by the lower pairwise interaction energies of the non-canonical interface from previous MD simulations^38^. Collectively, these non-canonical CypA-CA interfaces act as a secondary binding site, once CypA binds to the canonical CypA-binding loop. While we focused on the primary CypA binding site, since mutation and protonation of residue H219 does not affect the dynamics of the residues at the non-canonical interfaces, it is conceivable that CypA binding at these secondary sites may not be regulated by the H219Q mutation.

**Figure 6:**
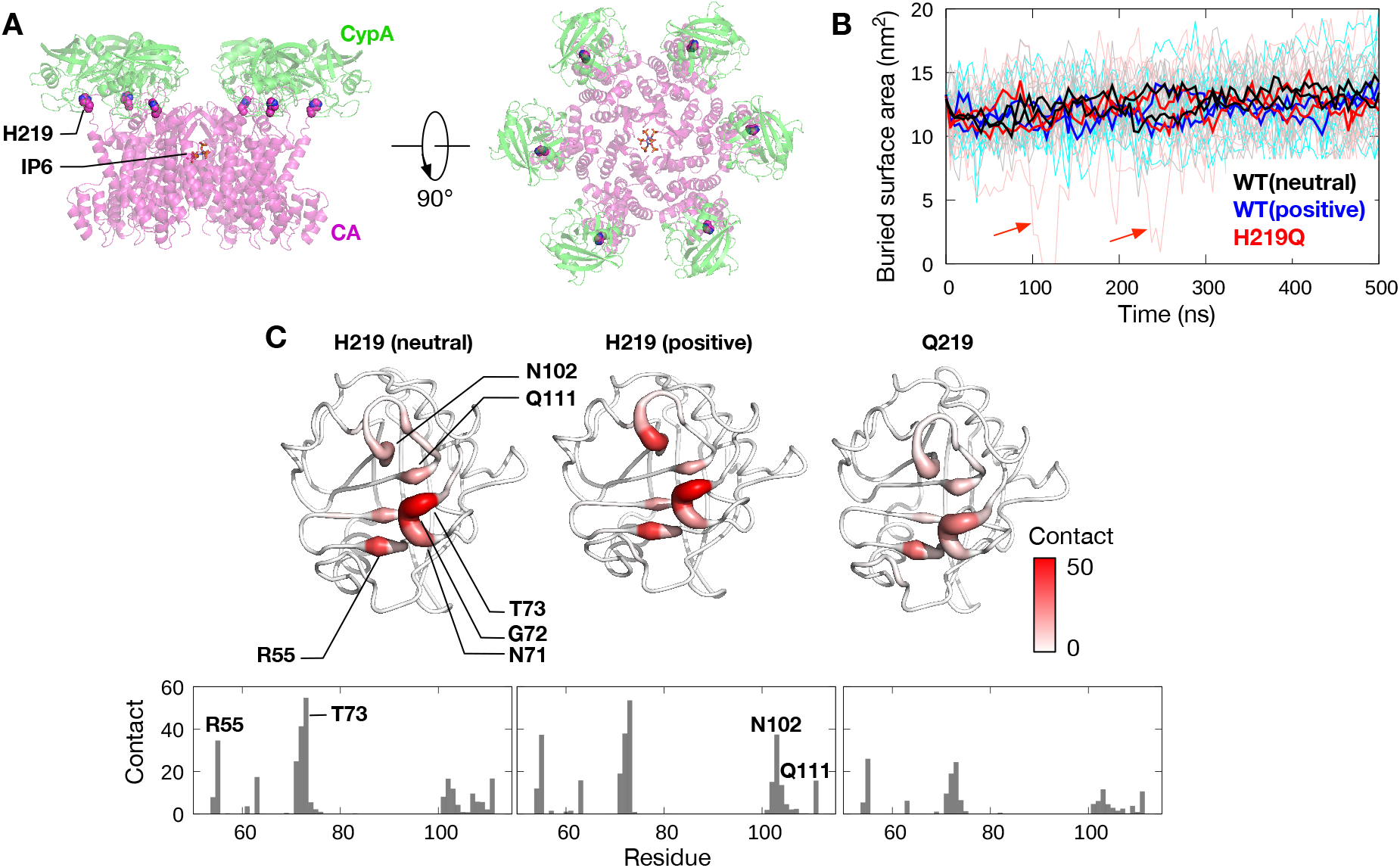
Atomistic simulations of mature CA with CypA. (A) Side and top views of mature CA hexamer (PDB: 6BHT) with each subunit bound to CypA (aligned using PDB: 1AK4). Residue H219 is shown in van der Waals representation, while IP6 bound in the center of the hexamer is shown in stick representation. (B) Buried surface area between CA and CypA throughout 500 ns simulations. Data taken from two independent simulations of CA with neutral H219 (black), protonated H219 (blue) and H219Q mutant (red). Thin lines indicate values from each of the six subunits, whilst the thick lines show the running averages. Red arrows show transient dissociation of CypA from CA in H219Q mutant simulations. (C) Residues on CypA that made contacts specifically with the side chains of neutral and protonated H219 (left and centre, respectively) from the WT CA simulations and Q219 (right) from the mutant CA simulations. CypA is shown in ribbon representation, the colour and thickness of which represent the percentage of contact made with CA during the simulations. A corresponding plot of per-residue contact percentages for each system is shown underneath, with key residues labelled. The cut-off distance used for contact analysis was 0.4 nm.

### Mutation in the CA hexameric interface stabilizes oligomerisation

Finally, another non-cleavage site mutation in the CA domain occurs at position 199, in which a glutamine residue is mutated to a histidine^20^. In immature Gag, this residue is located inside the CA domain pointing towards the internal cavity. Our CG simulations show that this residue did not form any major interactions with other CA residues as it was fully exposed to the solvent within the cavity of the CA. However, as CA undergoes large conformational changes during maturation, the Q199 residue forms a part of the hexameric interface of a mature CA domain (Figure 7A). This residue bridges the N-terminal domain of one CA subunit and the C-terminal domain of an adjacent subunit via polar interactions with residues such as Y301, K314 and Q311. We hypothesized that a mutation to histidine may alter the interactions between the two CA subdomains and therefore affect oligomerisation.

To test this hypothesis, we performed two repeat 500 ns atomistic simulations of mature CA for the WT and Q199H variants, the latter in either protonated or unprotonated states (Simulations 7, 13 and 14 in Table S2) (Figure 7B). We found that both neutral and charged histidine retained all major interactions as observed for glutamine in the WT protein; for example, the two most prevalent interacting partners from the neighbouring subunit were Y301 and L343, via polar and van der Waals interactions, respectively. Similar to the WT simulations, these residues were found to interact with H199 throughout most of the trajectories. However, the mutant simulations also showed additional contacts with some residues compared to WT. Thus, neutral H199 made significantly more frequent interactions with K314 and V313, while protonated H199 could form a salt bridge with E319. Protonation of H199 also modulates some interactions at the hexameric interface; for example, contact with R305 is noticeably reduced due to electrostatic repulsion, whilst intermittent contacts were observed with Q311 and E312, which explains the large standard deviations between the two repeat simulations. Pair interaction energies of both neutral and protonated H199 with surrounding residues from the neighbouring subunit are significantly more negative than that of Q199 (Figure 7C), indicating stronger interactions made by this histidine residue at the CA hexameric interface compared to the wild type glutamine. Our simulations therefore suggest that a histidine residue at position 199 potentially strengthens the binding interface between CA subunits to enhance the stability of the hexameric complex.

**Figure 7:**
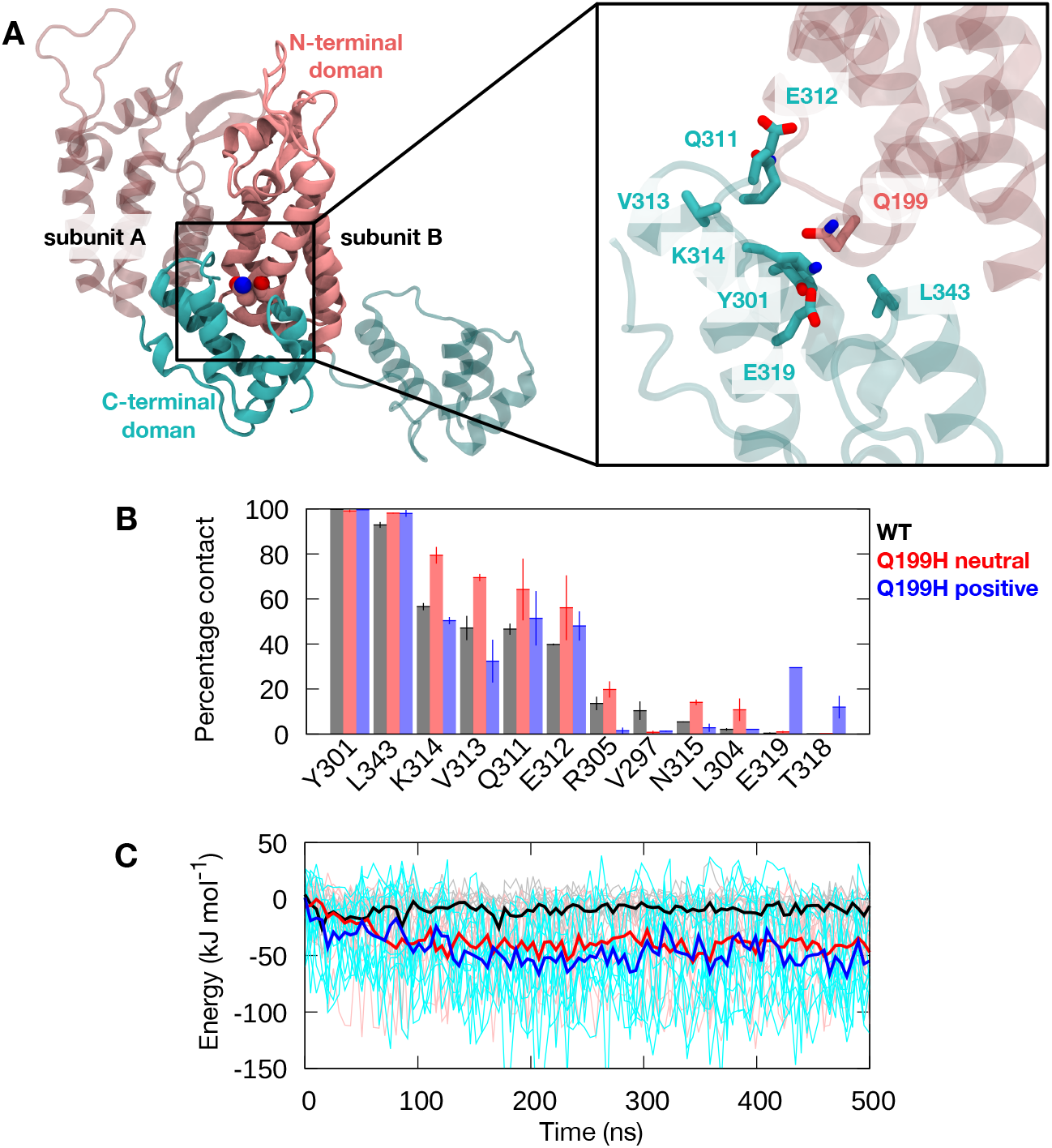
The role of the Q199H mutation in CA oligomerisation. (A) (Left) A snapshot of two subunits of the mature WT CA hexamer in cartoon representation. The Q199 residue bridging the N-terminal domain (pink) of one subunit and the C-terminal domain (cyan) of an adjacent subunit is shown in van der Waals representation. (Right) Enlarged image of the hexameric interface showing residues found within 0.4 nm of the Q199 residue. (B) Percentage contact made by residue Q199 (WT), neutral or positively charged H199 (mutants) with residues from adjacent CA subunits. Contacts are averaged over all six subunits and two independent simulations. Error bars indicate standard deviations between repeat simulations. The cut-off distance used for contact analysis was 0.4 nm. (C) Short-range Coulombic pairwise interaction energies between residue at position 199 on one subunit and surrounding residues on the adjacent subunit. Thin lines indicate values from each of the six subunits and two independent simulations, while thick lines show the running averages.

## Discussion

Given the possible benefits of drugging Gag together with the protease, we set out to study cleaved Gag structures using multiscale modelling and simulations of the HIV-1 Gag protein to understand the effect of clinically relevant mutations that occur outside of the cleavage sites. We built a model of the MA-CA-SP1 polyprotein complex in its multimeric state, which constitutes the largest cleavage product of the immature Gag protein. Our simulations of the membrane-bound complex suggest that the long linker connecting the MA and CA domains is flexible and may contract, enabling interactions between protease cleavage site residues and the C-terminal region of the MA domain, as well as several regions on the CA domain including the CypA binding loop. Two clinical mutations, G123E and H219Q, are found in these regions and alter the accessibility and electrostatic properties of the protease binding site. Mutations in the MA domain are concentrated on the predicted membrane binding surface and at least three of them – E12K, E40K and L75R – render the surface more positively charged compared to wild type Gag. Our CG simulations of these mutant variants show more pronounced interactions with PIP2 lipids, leading to the hypothesis that these mutations improve Gag targeting and anchoring to the plasma membrane during viral assembly. Atomic-resolution simulations of the mature Gag CA domain showed that the H219Q mutation may play a role in optimizing CypA interactions by controlling the conformational flexibility of the CypA binding loop and fine-tuning the CypA binding affinity. We also found that a mutation in the CA domain, Q199H, likely enhances oligomerisation by providing more stabilising interactions between adjacent subunits.

In the field of HIV and other related retroviruses, multiscale modelling and simulations have been utilised to study CA in different maturation states, providing important insights into their structural dynamics and interactions with drugs and host cell proteins^38,67-69^. For example, landmark microsecond-timescale atomic simulations of the entire HIV-1 capsid revealed important physicochemical properties, such as its electrostatics, dynamics, and water/ion permeability, as well as pointing towards long-range allosteric motions in the shell^35^. Simulations of the monomeric MA domain shed light on the molecular mechanism underlying membrane anchoring by the myristoyl group and PIP2 lipids^41^. Our study reports the first complete oligomeric model of the proteolytic product of HIV-1 Gag polyprotein, allowing the investigation of interactions between residues on the protease binding site and distant residues on the MA and CA domains, especially those from different subunits. Such inter-subunit “crosstalk” would be difficult to identify using individual crystal structures of mature Gag domains or a monomeric Gag model. Understanding structural changes that occur upon proteolysis could provide important clues on HIV-1 protease activity. For example, the MA-CA linker contraction observed in our simulations resulted in a significant decrease in solvent accessibility of the protease binding site, which helps to explain the lower cleavage rate of the MA-CA domain compared to the SP2-P6 domain^49^. From the perspective of anti-HIV drug development, targeting Gag proteolytic products in addition to the mature form, by virtue of identifying intermediate structural models, may allow for the discovery of Gag inhibitors that can inhibit multiple stages of maturation^14^.

Since accumulation of deleterious mutations could negatively impact virion survival, Gag polyprotein and protease co-compensate to outcompete protease inhibitors and restore viral fitness^14,15,70^. While HIV-1 protease acquires drug resistant mutations to evade PI activity, its ability to bind and subsequently cleave the Gag protein is impaired. Thus, Gag gains additional mutations to compensate for fitness defects to recover protease function. Structural studies show that associated mutations on cleavage sites do not restore interactions lost due to mutations on the protease, but rather establish novel interactions surrounding the site of protease mutations as well as induce conformational changes to drive better binding^18^. Apart from cleavage site mutations, mutations in more distant regions have been shown to contribute towards recovery from the reduced viral fitness caused by protease mutations^19,20^. Our study shows that these non-cleavage site mutations have far reaching implications outside of Gag proteolysis, from improving membrane binding to fine-tuning CypA interactions and stabilising the CA hexamer. A recent single genome sequencing study revealed that, indeed, most Gag-protease correlated mutations occur outside of the Gag cleavage site with the viral MA and CA domains representing the largest subsets of the mutations^70^. Interestingly, a high concentration of compensatory mutations was uncovered within the globular domain of MA (which binds to the plasma membrane) and the CypA binding loop of CA, further corroborating our simulation results. Mutations and deletions of residues on the C-terminal region of the MA domain, which have direct contact with cleavage site residues based on our simulations, have also been recently shown to confer resistance to PI lopinavir^71^. Our results also highlight that the residues on the CypA binding loop can interact directly with MA-CA cleavage site residues, thus potentially influencing their local electrostatics and surface accessibility, although determining the exact interactions made between these residues and subsequent molecular effects would require further studies. Additionally, PS lipids are known to be exposed on the outer leaflet of HIV-1 membrane and may facilitate HIV-1 entry into the host cells^72,73^, and it is possible that the highly basic surface generated by the non-cleavage site mutations could attract anionic PS lipids, similarly to the PIP2 enrichment around the MA domain observed here. While our study assumes that these non-cleavage site mutations do not have any large impact on the overall structure of Gag, it is possible that these point mutations could result in local structural changes. Further structural and computational studies would thus be of interest to directly elucidate the impact of these mutations on Gag structure. It is worth noting that compensatory non-cleavage site mutations also occur in the NC and P6 domains. Gatanaga et al. discovered five such mutations with clinical importance^20^, and many clinical drug resistant mutations can occur within the first replication cycle^7^. While modelling these domains is beyond the scope of the current study, it is certainly worth exploring in the future, specifically regarding how these mutations affect interactions with the viral RNA genome. Overall, our data have revealed new insights into the role of Gag non-cleavage site mutations on HIV-1 viral fitness. Given that HIV-1 is one of the fastest mutating RNA viruses, an in-depth comprehension of emerging mutations will aid in the general understanding of viral drug resistance, as well as the emergence of novel viruses that are able to cross species during zoonotic infections.

## Materials and Methods

### Model building and system setup

The model of HIV-1 Gag MA-CA-SP1 was built using two structural templates: MA trimers (PDB: 1HIW)^42^ and the immature CA hexamer (PDB: 5L93)^43^. To the best of our knowledge, the former represents the only available crystal structure of the MA protein in its trimeric form, whilst the latter represents the highest resolution structure of a complete immature CA hexamer. Six copies of MA trimers were arranged based on EM images showing that they organize as hexamer-of-trimers on a model membrane bilayer^44^. The CA hexamer was placed such that the N-termini of its subunits align with the C-termini of the central subunits of the MA trimers. The MA and CA domains are connected by a flexible, disordered linker as shown by solution NMR studies^45,46^. Modeller version 9.21^74^ was used to build the loops connecting the MA trimers and CA hexamer (Figure S1) using the discreet optimized protein energy (DOPE)-based loop modelling protocol, and the model with the lowest DOPE score was chosen^75^. Stereochemical assessment using Ramachandran analysis^76^ showed only one outlier residue, confirming that the model was structurally reasonable. The atomistic MA-CA-SP1 structure was subsequently converted to CG representation using the MARTINI 2.2 force field^77^ and the elastic network model, ElNeDyn^78^, was imposed to maintain the integrity of the secondary and tertiary structure. This CG model was validated by comparing the MA-CA domain dynamics to all-atom simulations of a monomeric MA-CA-SP1 protein (more details below). The N-terminal glycine residue on each MA subunit was myristoylated based on parameters from Charlier et al.^41^ Three mutant variants were generated using PyMOL^79^ based on non-cleavage site mutations published by Gatanaga et al.^20^ The wild-type residues were substituted with the mutant versions using the PyMOL mutagenesis tool, assuming these point mutations would not result in any large structural deviations from wild-type Gag.

A 30 x 30 nm^2^ patch representing the HIV-1 membrane model was built using the CHARMM-GUI Martini Maker Bilayer Builder^80,81^ and the lipid composition was based on a previous HIV-1 lipidomic study^47^. This membrane was asymmetric: the upper leaflet was made of 10% phosphatidyl choline (PC), 10% phosphatidyl ethanol amine (PE), 30% sphingomyelin, and 50% cholesterol, whereas the lower leaflet was made of 10% PC, 20% PE, 15% phosphatidyl serine (PS), 5% phosphatidylinositol-4,5-bisphosphate (PIP2), and 50% cholesterol. The HIV-1 membrane is enriched in saturated lipid species especially for PC; we therefore modelled the lipid tails of PC as 1,2-dipalmitoyl (DP), whereas the rest of the lipids were modelled with 1-palmitoyl-2-oleoyl (PO) lipid tails. Following minimization and equilibration procedures according to the CHARMM-GUI, the model membrane was further simulated for 1 μs to allow better mixing of the lipid components prior to simulations in the presence of protein.

### Molecular dynamics simulations

The HIV-1 Gag MA-CA-SP1 CG model was placed underneath the membrane such that the myristoylated N-termini were inserted into lower leaflet of the membrane. Energy minimization was then performed using the steepest descent method to remove any overlapping beads. The system was solvated with standard MARTINI water molecules and neutralized with 0.15 M NaCl ions. Further energy minimization was performed. The system was then equilibrated for 10 ns whereby positional restraints with a force constant of 1000 kJ mol^-1^ nm^-2^ were imposed on all of the backbone atoms of the protein. Temperature coupling using the V-rescale thermostat with a time constant of 1 ps^82^ was applied to maintain the temperature at 310 K. Semi-isotropic pressure coupling using the Berendsen barostat with a time constant of 5 ps was utilized to maintain the pressure of the system at 1 atm. Electrostatics were calculated using the reaction field method. The der Waals interactions were computed using a potential shift Verlet scheme. The short-range cut-offs for both of these were set to 1.1 nm. After the equilibration simulations, four independent production runs of 15 μs were performed using different distributions of initial velocities. The same protocols used for the equilibration simulation were used for the production runs in the absence of position restraints, except for the pressure coupling in which the Parrinello-Rahman barostat was used with a coupling constant of 12.0 ps^83^.

Atomistic MD simulations were performed to understand the role of the H219Q mutations on CypA binding and Q199H on CA oligomerisation. The structure of the mature HIV-1 CA hexamer in complex with inositol IP6 (PDB: 6BHT)^40^ was aligned to that of the CA N-terminal domain bound to CypA (PDB: 1AK4)^55^. This alignment generated a structure of the CA hexamer bound to six copies of CypA (Figure 6A). Mutations in the CA were performed using PyMOL. The PROPKA3 programme^58^ was used to calculate the pKa of H219 (Table S3). The proteins were parameterized using the CHARMM36 force field^84^, whereas the CHARMM-GUI Ligand Reader & Modeller^85^ was used to generate the parameters for the IP6 molecule. The complex was solvated with TIP3P water molecules and 0.15 M NaCl was added to neutralize the system. The steepest descent minimization protocol was used to remove overlapping atoms. A short 100 ps equilibration simulation with position restraints (force constant of 1000 kJ mol^-1^ nm^-2^) applied to all heavy atoms of the protein was conducted. The temperature of the system was maintained at 310 K using the Nosé-Hoover thermostat with a time constant of 1.0 ps^86^,^87^. The pressure was kept at 1 atm using an isotropic coupling to the Parrinello-Rahman barostat with a time constant of 5.0 ps^83^. Electrostatic interactions were measured using the smooth particle mesh Ewald (PME) method with a real-space cut off of 1.2 nm^88^. The van der Waals interactions were calculated using the force switch smoothing function applied between 1.0 and 1.2 nm, and truncated at 1.2 nm. An integration time step of 2 fs was used with all covalent bonds involving hydrogens constrained using the LINCS algorithm^89^. After equilibration, the positional restraints were removed and two independent 500 ns production runs were conducted using the same setup for each of the WT and mutant variants starting with different initial velocities. To assess whether the systems were properly equilibrated, we calculated the RMSD of the Cα atoms of CA (Simulation 7-14 in Table S2) and CypA (Simulation 10-12 in Table S2) for all six chains of the hexamer (Figure S10). We found that in most cases the RMSD reached a plateau after around 100 ns.

Additional atomistic and CG simulations were performed to verify the conformation of the MA-CA linker. A single subunit of the HIV-1 Gag MA-CA-SP1 model was placed in solution with 0.15 M NaCl ions. Positional restraints with a force constant of 1000 kJ mol^-1^ nm^-2^ were applied to the backbone atoms of membrane binding residues (residue 2-53 and 72-90) to mimic membrane binding. Four independent 500 ns simulations were performed for both atomistic and CG systems using the same protocols described above.

To refine interactions between the linker region and mutant residues observed in the CG simulations of the MA-CA-SP1 hexamer, we converted the structure of the hexamer from the final frame of one of the simulations to atomistic representation using the CHARMM-GUI All-Atom Converter. The back-mapping protocol was performed using a flexible geometric approach via the *backward.py* program^50^. In this approach, the atomic particles are mapped to the weighted average of the CG beads, followed by geometrical corrections whereby the atomic particles are repositioned (when necessary) to reconstruct chiral centres and double bond configurations. Force field-based relaxation consisting of a series of energy minimizations and position restrained MD simulations was then conducted. A single subunit of the MA-CA-SP1 was subsequently used to refine interactions with the G123E mutant, whereas two adjacent subunits were used for interactions with the H219Q mutant. The protein was inserted into a box of water with 0.15 M NaCl ions. Positional restraints with a force constant of 1000 kJ mol^-1^ nm^-2^ were imposed on the backbone atoms of residues outside of the MA-CA linker and CypA binding loop (residue 2-117, 147-215 and 233-376). For each system, three independent 200 ns simulations were performed using the parameters described above.

All simulations were performed using GROMACS 2018^90^. The list of simulations performed is provided in Table S2. Analysis of the electrostatic surface charge of the MA domain was performed in PyMOL using the APBS plugin^91^. Cluster analysis of the CypA binding loop (residue 216-232) was performed on structures generated from simulation trajectories using the GROMACS *gmx cluster* tool. The clustering algorithm described by Daura et al.^92^ was applied using a cutoff of 0.3 nm.

## Acknowledgments

This work was supported by BII and EDDC of A*STAR. Simulations were performed on the petascale computer cluster ASPIRE-1 at the National Supercomputing Centre of Singapore (NSCC) and the A*STAR Computational Resource Centre (A*CRC).

