## Supplementary for "The Impact of Gag Non-Cleavage Site Mutations on HIV-1 Viral Fitness from Integrative Modelling and Simulations" (from another version of this preprint: version 1, 2020-07-05)

**Table S1: List of non-cleavage site mutant variants (Gatanaga et al., 2002)**

| Mutant variant | Mutations |
| --- | --- |
| 1 | V35I-L75R-H219Q |
| 2 | L75R-H219Q-R409K-L449F-E468K |
| 3 | E12K-V35I-L75R-H219Q-V390D-R409K-L449F-E468K |
| 4 | H219Q-V390D |
| 5 | H219Q-V390D-R409K-L449F |
| 6 | V35I-E40K-G123E-H219Q |
| 7 | E12K-E40K-G123E-Q199H-H219Q-R409K-G412D-L449F-E468K |

**Table S2: List of simulations**

| Protein | Variant | Resolution | Simulation time |
| --- | --- | --- | --- |
| MA-CA-SP1 | WT | Coarse-grained | 4 X 15 µs |
| MA-CA-SP1 | Mutant 3 | Coarse-grained | 4 X 15 µs |
| MA-CA-SP1 | Mutant 6 | Coarse-grained | 4 X 15 µs |
| MA-CA-SP1 | Mutant 7 | Coarse-grained | 4 X 15 µs |
| CA | WT (H219 neutral) | Atomistic | 2 X 500 ns |
| CA | WT (H219 protonated) | Atomistic | 2 X 500 ns |
| CA | H219Q | Atomistic | 2 X 500 ns |
| CA + CypA | WT (H219 neutral) | Atomistic | 2 X 500 ns |
| CA + CypA | WT (H219 protonated) | Atomistic | 2 X 500 ns |
| CA + CypA | H219Q | Atomistic | 2 X 500 ns |
| CA | Q199H (neutral) | Atomistic | 2 X 500 ns |
| CA | Q199H (protonated) | Atomistic | 2 X 500 ns |
| MA-CA-SP1 monomer | WT | Atomistic | 4 X 500 ns |
| MA-CA-SP1 monomer | WT | Coarse-grained | 4 X 500 ns |

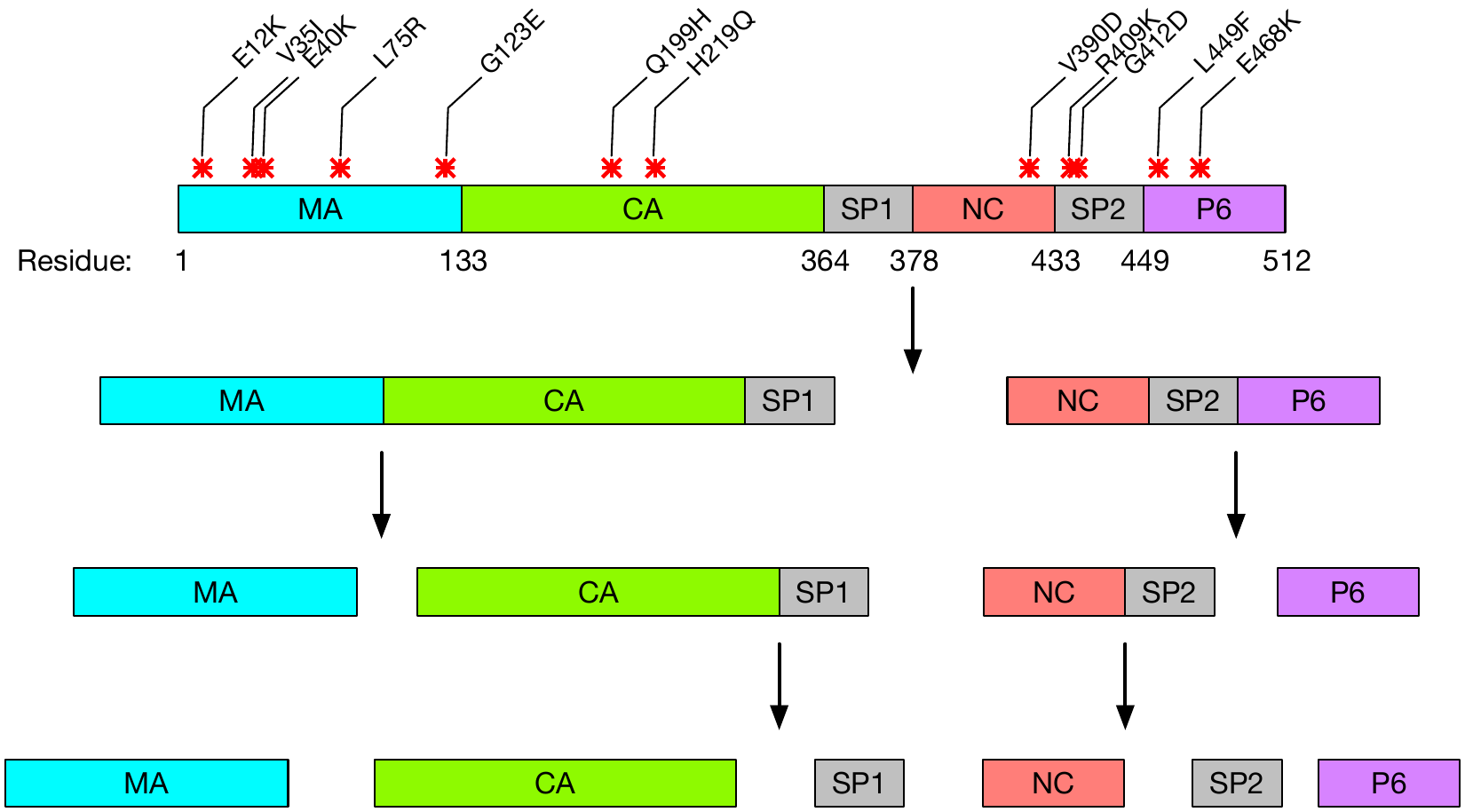

**Figure S1: A schematic of the sequential proteolysis of HIV-1 Gag polyprotein.** The approximate positions of non-cleavage site mutations discovered by Gatanaga et al. (Table S1) are marked with an asterisk.

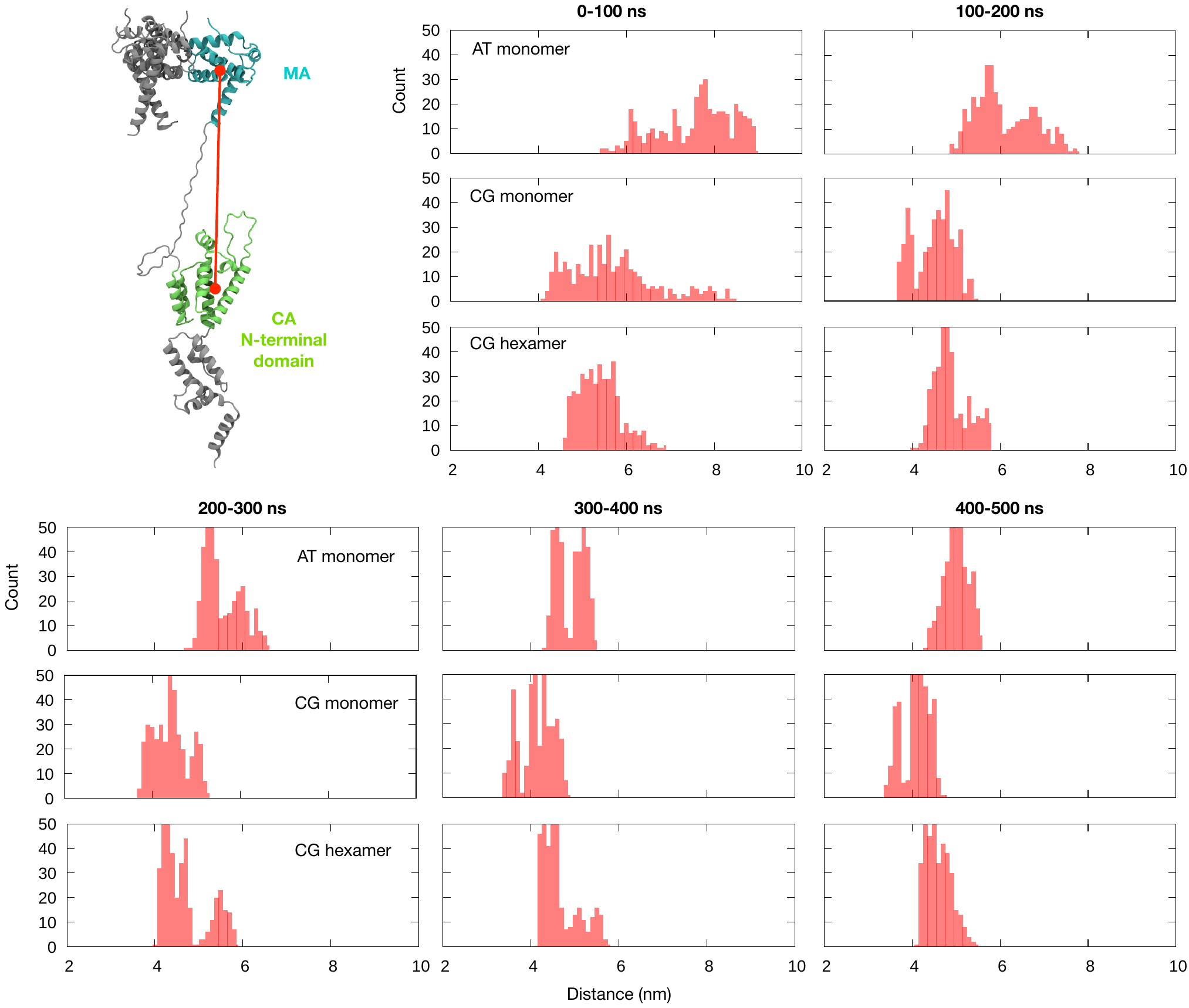

**Figure S2: Comparison of domain dynamics between atomistic (AT) and coarse-grained (CG) simulations during MA-CA linker contraction.** The distributions of distance between the centers of mass of the MA domain (residue 1 to 114) and the N-terminal region of the CA domain (residue 150-275) from four independent 500 ns simulations are shown, presented in 100 ns intervals. Data were taken from simulations of MA-CA-SP1 monomer (top for atomistic and middle for CG) or hexamer (bottom).

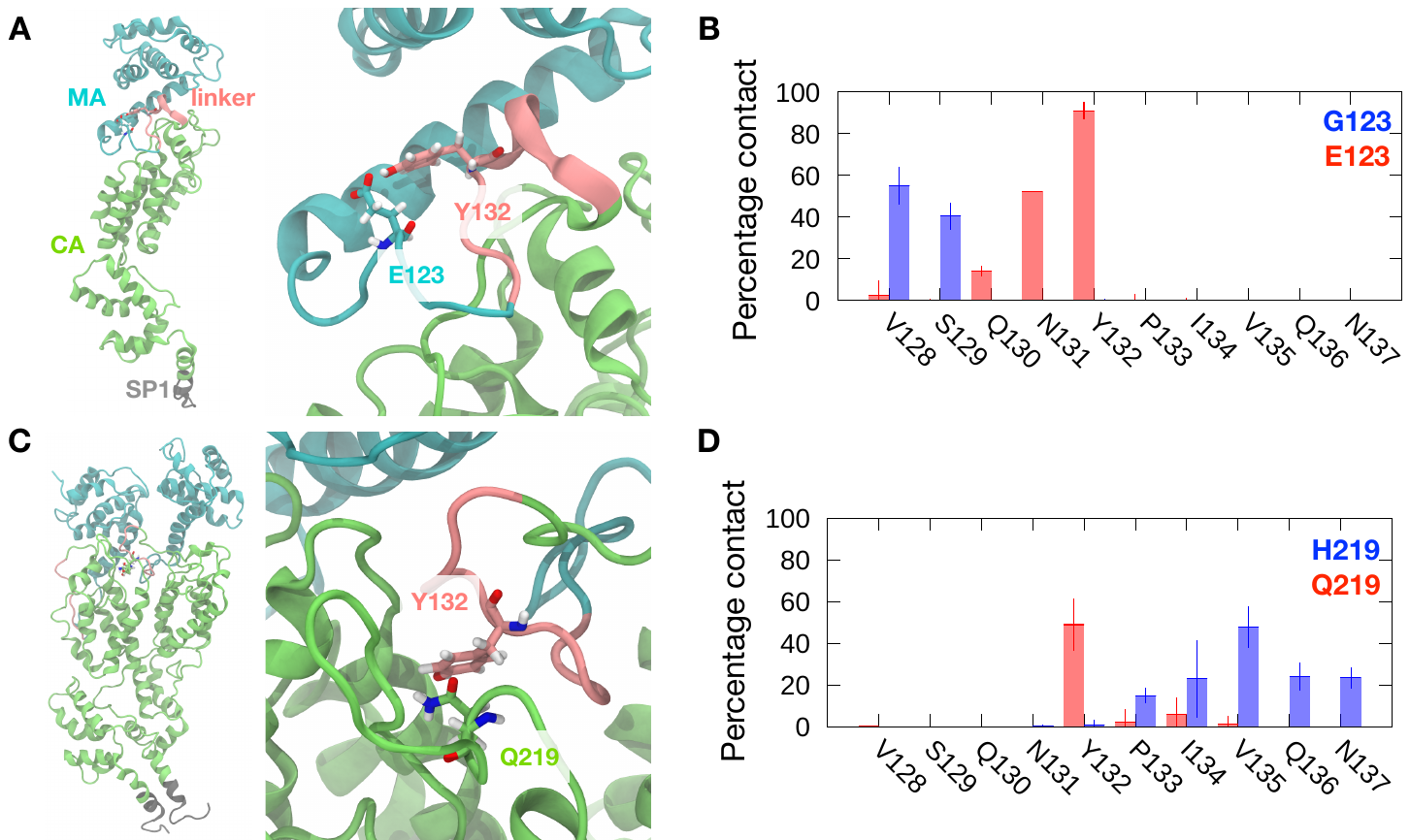

**Figure S3: Interactions of the MA-CA linker with mutant residues from atomistic simulations**. (A) Snapshots at the end of an atomistic simulation of G123E mutant using structures back-mapped from CG simulations. Full system shown on the left and enlarged image of E123 is shown on the right. A single MA-CA-SP1 subunit was used for this simulation. (B) Contact analysis performed between residue at position 123 (glycine in WT and glutamate in mutant) with residues on the MA-CA linker. Average values from three independent 200 ns simulations are shown, and error bars indicate standard deviations between replicates. (C) Snapshots at the end an atomistic simulation of the H219Q mutant using structures back-mapped from CG simulations. Full system is shown on the left and enlarged image of Q219 is shown on the right. Two adjacent MA-CA-SP1 subunits were used for this simulation. (D) A similar contact analysis performed on residue at position 219 (histidine in WT and glutamine in mutant).

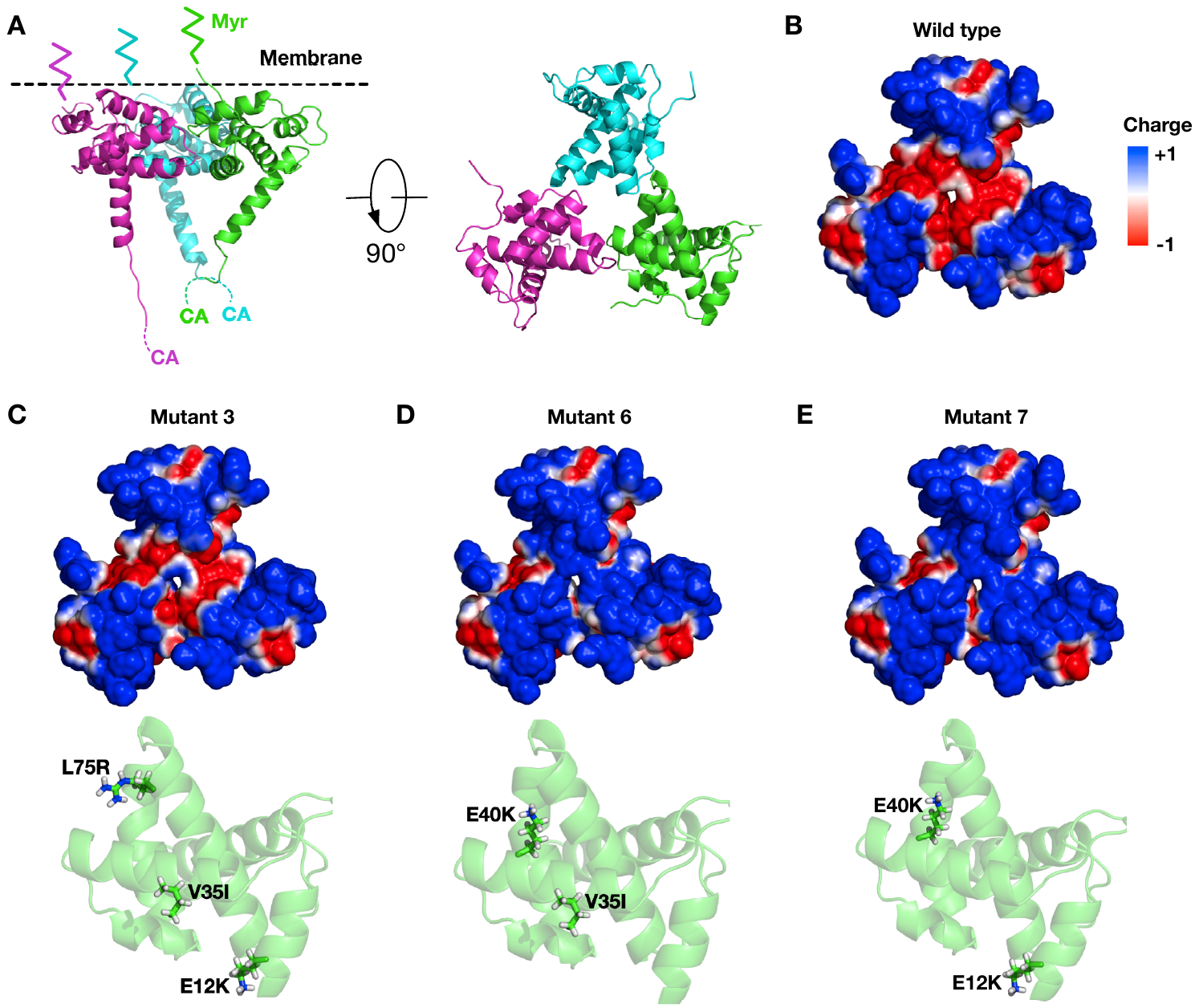

**Figure S4: Electrostatic surface charge of MA domain.** (A) The MA trimer as viewed along the plane of the membrane (left) and its membrane binding surface (right). (B) The electrostatic surface charge of the membrane binding surface of wild type MA. (C-E) The same analysis performed on mutants 3, 6 and 7. The positions of mutated residues are mapped onto one MA subunit and are shown underneath.
